# Injectable Myocardial Matrix Hydrogel Mitigates Negative Left Ventricular Remodeling in a Chronic Myocardial Infarction Model

**DOI:** 10.1101/2020.07.31.231449

**Authors:** Miranda D. Diaz, Elaine Tran, Jean W. Wassenaar, Martin Spang, Roberto Gaetani, Colin G. Luo, Rebecca Braden, Ryan C. Hill, Kirk C. Hansen, Anthony N. DeMaria, Karen L. Christman

**Affiliations:** Department of Bioengineering, University of California, San Diego; Sanford Consortium for Regenerative Medicine; Department of Molecular Medicine, Sapienza University of Rome; Department of Medicine, University of California, San Diego, La Jolla, California; Department of Biochemistry and Molecular Genetics, University of Colorado, Anschutz Medical Campus

**Keywords:** Heart failure, biomaterials, gene expression, chronic inflammation

## Abstract

A first-in-man clinical study on a myocardial-derived decellularized extracellular matrix (ECM) hydrogel yielded evidence for potential efficacy in ischemic heart failure (HF) patients. However, little is understood about the mechanism of action in chronic myocardial infarction (MI). In this study we investigated efficacy and mechanism by which the myocardial matrix hydrogel can mitigate negative left ventricular (LV) remodeling in a chronic model of MI. Assessment of cardiac function via magnetic resonance imaging (MRI) demonstrated preservation of LV volumes and apical wall thickening. Differential gene expression analyses showed the matrix is able to prevent worsening HF in a small animal chronic MI model through modulation of the immune response, downregulation of pathways involved in HF progression and fibrosis, and upregulation of genes important for cardiac muscle contraction.

## Introduction

More than 6 million Americans are currently living with heart failure (HF), and this number is continuing to rise as the average global age increases and therapy of causative conditions improves (1). Currently, there are various injectable biomaterials are being investigated in pre-clinical models mainly for treatment of acute or sub-acute MI (2, 3). However, little has been done in investigating the efficacy of injectable biomaterial therapeutics in pre-clinical chronic MI or ischemic HF models. We previously developed a decellularized injectable extracellular matrix (ECM) hydrogel derived from porcine LV myocardial tissue as a tissue engineering approach for treatment after MI (4). While the macrostructure of the ECM is broken down in processing, the myocardial matrix hydrogel retains the nanostructure, peptides, and glycosaminoglycans (sGAGs) found in native myocardium and can be delivered minimally invasively via transendocardial delivery. Sub-acute delivery of the hydrogel 1-2 weeks post-MI was previously shown to mitigate negative LV remodeling and improve cardiac function in small and large animal models with sub-acute delivery of the material 1-2 weeks post-MI (4–6). In a rat model of sub-acute MI, involvement of specific pro-repair pathways was found in response to delivery of the hydrogel, including modulation of the inflammatory response, altered metabolism, upregulation of genes involved in neovascularization, and downregulation of genes involved in hypertrophy, fibrosis, and apoptosis (6). The myocardial matrix hydrogel, manufactured as VentriGel, showed safety in a first-in-man phase I clinical trial with a decellularized ECM hydrogel therapeutic (7). While the study was not designed to evaluate efficacy, magnetic resonance imaging (MRI) analysis showed that reduction in LV volumes was observed predominantly in patients more than one year post-MI suggesting that the material can be effective in chronic MI, however, little is understood about the mechanism of action in this patient population. Therefore, in this study we aimed to investigate the efficacy and mechanism of action of the myocardial matrix hydrogel in preventing the progression of negative LV remodeling and dysfunction. Understanding the differences in mechanism of action of the material between the sub-acute and chronic post-MI environments could guide the next phases of clinical testing of the hydrogel as well as influence the understanding of therapeutic targets for the design of future MI therapeutics.

## Materials and Methods

### Myocardial Matrix Preparation and Characterization

Myocardial matrix was prepared from porcine left ventricular myocardial tissue and characterized as previously outlined (4, 8). Briefly, the tissue was chopped, decellularized in sodium dodecyl sulfate (SDS) detergent, lyophilized, milled into a fine powder, partially enzymatically digested with pepsin in HCl, adjusted for pH and salts, brought to a final concentration of 6 mg/mL, and finally aliquoted and lyophilized for storage at −80°C. When preparing for injection, lyophilized aliquots were resuspended in sterile water approximately 30 minutes prior. In addition to our standard published assays for quality control, we also performed ECM targeted, QconCAT proteomics on the decellularized ECM as previously described (8–10).

### Surgical Procedures

All procedures in this study were performed in accordance with the guidelines established by the Committee on Animal Research at the University of California, San Diego and the Association for the Assessment and Accreditation of Laboratory Animal Care. All animals used in the study were adult female Sprague Dawley rats (225-250g). To induce MI, all animals underwent occlusion-reperfusion surgery to occlude the left main artery for 35 minutes as previously described (4). At 8 weeks post-MI, animals were arbitrarily assigned to injection of 75 μL of either matrix or saline directly into the infarct via subxiphoid access as previously described (4, 11).

### Magnetic Resonance Imaging (MRI)

Cardiac Cine MR images were acquired using an 11.7T Bruker MRI System by Molecular Imaging Inc. at the Sanford Consortium for Regenerative Medicine. Rats were anesthetized using isoflurane in oxygen during imaging. Respiratory and ECG-gated, cine sequences were acquired over contiguous heart axial slices. The following parameters were used: repetition time = 20 ms, echo time = 1.18 ms, flip angle = 30°, field of view = 40 mm2, data matrix size = 200 x 200. Eight or nine 1.5 mm, axial image slices were acquired with a total of 15 cine frames per image slice. ImageJ (NIH) was used to outline the endocardial surface at end diastole and end systole for each slice, defined as the minimum and maximum LV lumen area, respectively. Simpson’s method was used to calculate the end-diastolic volume (EDV) and end-systolic volume (ESV). Ejection fraction (EF) was calculated as [(EDV-ESV)/EDV] x 100. The myocardium was defined as the area between the epicardium and endocardium. Myocardial area was analyzed at EDV and ESV to assess wall thickening by myocardial area (12, 13). Cardiac wall thickening was calculated as [(EDV myocardial area-ESV myocardial area)/EDV myocardial area] from cross sectional MR images the of basal, mid, and apical slices. Rats underwent baseline MRI two-days prior to injection of matrix or saline (8 weeks post-MI). Animals that did not have an EF at least one standard deviation below healthy values (< 68%) were excluded from the study. A total of 48 rats underwent the MI procedure, 16 died as a result of the MI, and 5 were excluded based on EF criteria leaving n=8 for saline and n=6 for matrix for final analysis. At twelve-weeks post-MI (four-weeks post-injection), rats were imaged and their cardiac function assessed again for posttreatment evaluation. All MRI acquisition and analyses were performed by investigators blinded to the treatment groups.

### Tissue Processing

Another set of animals (n=6 each group) were euthanized at 1 week post-injection and their hearts were cut into 7 slices using a stainless steel rat heart slicer matrix (Zivic Instruments) with 1.0 mm coronal spacing. The infarct region was isolated from even slices of tissue and flash frozen in liquid nitrogen to preserve RNA. The remaining slices were fresh frozen in TissueTek OCT™ freezing medium. For animals that were euthanized 5 weeks post-injection (~1 week post final MR imaging), hearts were fresh frozen in TissueTek OCT™. Once frozen, the heart tissue was cryosectioned into 10 um sections.

### Cardiac fibroblasts production of MMPs

Cardiac fibroblasts were collected as the adherent fraction to tissue culture plastic at the end of cardiomyocyte isolation as described previously(14). Fibroblasts were cultured in 1 g/L glucose DMEM with 10% FBS and 1% P/S. Cells were passaged with trypsin once 90% confluency is reached and plated on either collagen or myocardial matrix hydrogels formed in 48-well plates at 120,000 cells/well (n = 6). The elution and digestion fractions of the hydrogel were generated as follows. After overnight gelation of the matrix, PBS (900 μL) was added on top of the 100 μL hydrogels and collected 24 hours later as the elution fraction. After the elution fraction was extracted, the solid portion of the hydrogels was either digested with MMP2 and MMP9 or incubated with the MMP activation buffer alone. The samples were centrifuged after 24 hours and the supernatant was isolated as the digestion fraction. Fibroblasts were then also seeded in 6-well plates at 500,000 cells/well and cultured in serum free media and the degradation products or their respective enzyme only controls at the 1:100 dilution (n = 3). Cells were cultured for 48 hours after which a sample of media from each well was collected. Media were mixed with equal volume of Tris-Glycine SDS Sample Buffer (Life Technologies) and analyzed using Novex Zymogam 10% Gelatin Gels (Life Technologies) per manufacturer instructions. Amersham ECL Full-Range Rainbow Molecular Weight Markers were used as the protein ladder. Gels underwent electrophoresis using the XCell SureLock Mini-Cell for 90 minutes under constant 125 V. Gels were developed using renaturing buffer and developing buffers, then visualized with Imperial Protein Stain. Band intensity was quantified by ImageJ (NIH) Gel.

### Histology

Slides were sectioned starting ~200 *μm* from the apex with 16 locations taken at 8 slides per location with 150 *μm* spacing between each location to span the ventricle. These slides were stained with H&E or Masson’s trichrome, mounted with Permount (Fisher Chemical) and scanned at 20x using an Aperio Scan Scope CS2 slide scanner (Leica Biosystems). H&E staining was used to identify 5 representative slides for each animal that were evenly spaced to span the infarct in the ventricle. Trichrome stained slides were used to assess fibrosis at five-weeks post-injection using five evenly-spaced representative slides throughout the infarct scanned at 20× magnification using an Aperio ScanScope CS2 slide scanner (Leica). Non-nuclear blue staining was measured using the ‘Positive Pixel Count V9’ algorithm within ImageScope (Aperio) software to determine the collagen content.

### NanoString Multiplex Gene Expression Analysis

RNA was isolated using the Qiagen RNeasy Fibrous Tissue Mini Kit and the QIAcube Connect nucleic acid isolation robot. For screening of various pathways of interest, RNA samples were analyzed by NanoString nCounter® MAX Analysis System (15, 16) with nCounter® custom cardiac codeset (Rat) allowing for multiplexed assessment of 380 genes (see Supplemental Appendix Table S4 for full gene list). Samples were processed according to manufacturer instructions. In brief, RNA sample concentrations were measured on a Qubit 3.0 Fluorometer with a Qubit™ RNA HS Assay kit. 70 μL of hybridization buffer was mixed with Immunology Panel Reporter CodeSet solution, and 8 μL of this master mix was mixed in a separate reaction vessel with 50-100 ng of RNA per tissue sample and RNA-free water up to 13 μL total. 2 μL of Capture ProbeSet was added to each vessel, mixed and placed on a thermocycler at 65°C for 16-48 hours before being maintained at 4°C for less than 24 hours. NanoString nCounter Prep Station performed automated fluidic sample processing to purify and immobilize hybridized sample to the cartridge surface. Digital barcode reads were analyzed by NanoString nCounter® Digital Analyzer. Results were analyzed by manufacturer nSolver^™^ Analysis Software 4.0 and custom R scripts. Outliers were detected and excluded based on NanoString’s methods for outlier detection.

### Statistical Analysis

*In vivo* functional and histological comparisons between matrix and saline treatment groups were made using a paired t-test when comparing baseline to post-injection and student’s t-test when comparing matrix treated to saline control group. Significance was accepted at p < 0.05. Data are reported as mean ± standard error of mean. For *in vitro* experiments, student’s t-test or one-way ANOVA was performed followed by Tukey’s post-hoc testing, where necessary. Gene expression normalization and differential expression was analyzed by the NanoStringDiff package with significance at a pvalue < 0.05 (17). Heatmap was displayed with the pheatmap package ([Software] KRpPh). Gene enrichment analysis was performed with the clusterprofiler package (18) and KEGG pathway images created with the Pathview package (19).

## Results

### Decellularized ECM Characterization

The myocardial matrix hydrogel was made using previously described decellularization methods and standard material characterization analyses were performed for batch quality control (8, 20). Here we also identified the specific protein components that remain post-decellularization using ECM-targeted Quantitative conCATamers (QconCAT) and a Liquid Chromatography - Selected Reaction Monitoring (LC-SRM) analysis. Out of 83 distinct peptides, which uniquely represent ECM and ECM-associated proteins, 28 were detected (Figure 1, Supplemental Table 1). We also performed global proteomics analysis using data dependent LC-mass spectrometry (MS)/MS analysis (Supplemental Table 2) since a few ECM proteins, such as collagen 3 alpha-1 chain, are not covered with the QconCAT peptides. The decellularized ECM is comprised mostly of fibrillar collagen (types I and III), but also includes numerous other ECM proteins and proteoglycans. The low percentage of cytoskeletal protein contribution (ACT and MYH) indicates the efficiency of the decellularization process (Figure 1, Supplemental Table 1).

**Figure 1.**
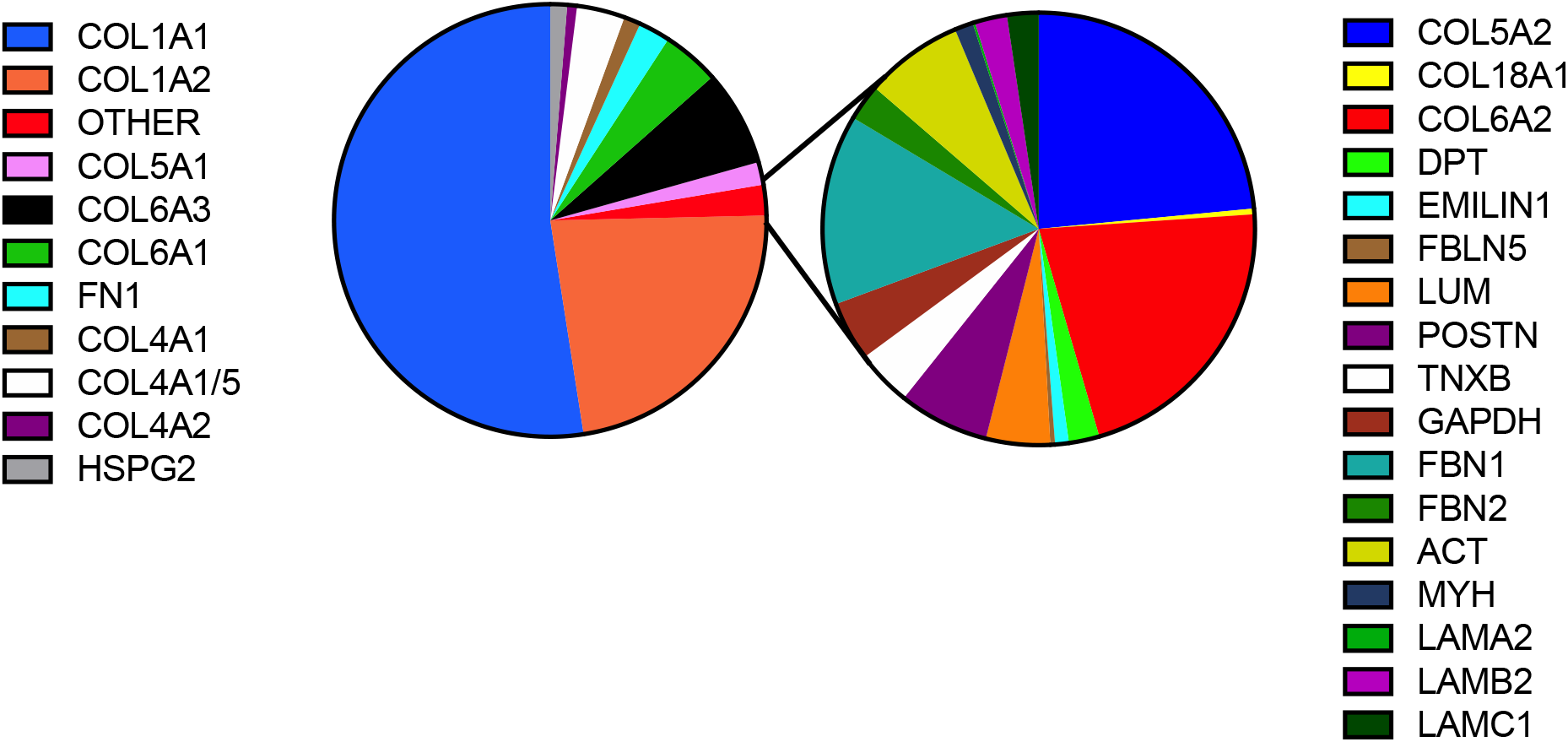
QconCat quantification of myocardial matrix hydrogel. Pie chart represents the average percentage of ECM, ECM associated, and cellular contaminant protein content as determine using targeted ECM proteomics.

### Effect of Myocardial Matrix Hydrogel on Negative LV Remodeling

While a phase 1 clinical trial showed evidence of potential efficacy for the matrix therapeutic in those treated more than 1 year post-MI, we wanted to confirm the effects on cardiac function in a chronic MI model (7). Animals were examined 8 weeks after MI to allow HF to develop and baseline MR images were taken 1-2 days pre-injection and 4 weeks post injection of either matrix (n=6) or saline (n=8). Comparing the images of matrix injected animals at baseline vs. post-injection, there were no significant changes in LV end systolic (131.2 mm^3^ vs. 135.1 mm^3^) and end diastolic volumes (340.9 mm^3^ vs. 367.9 mm^3^) (ESV/EDV) for the matrix treated group (n=6) (Figure 2c, d). The saline control group, however, shows significant increases from baseline to post-injection in ESV (148.9 mm^3^ vs. 174.8 mm^3^) and EDV (335.2 mm^3^ vs. 427.9 mm^3^) (n=7, p<0.05) (Figure2 c,d). Measurement of changes in myocardial wall thickening in the apical segment indicated that there was significantly reduced LV apical wall thickening in the saline control group (n=7) compared to matrix treated animals (Figure 2e). We previously found changes in fibrosis in a sub-acute MI model, and therefore evaluated infarct fibrosis 4 weeks after injection in the chronic MI model (20). Analysis shows trends (p=0.07) for reduced percentage of fibrosis in the infarct region of matrix injected hearts compared to the saline control group (Figure 3b).

**Figure 2:**
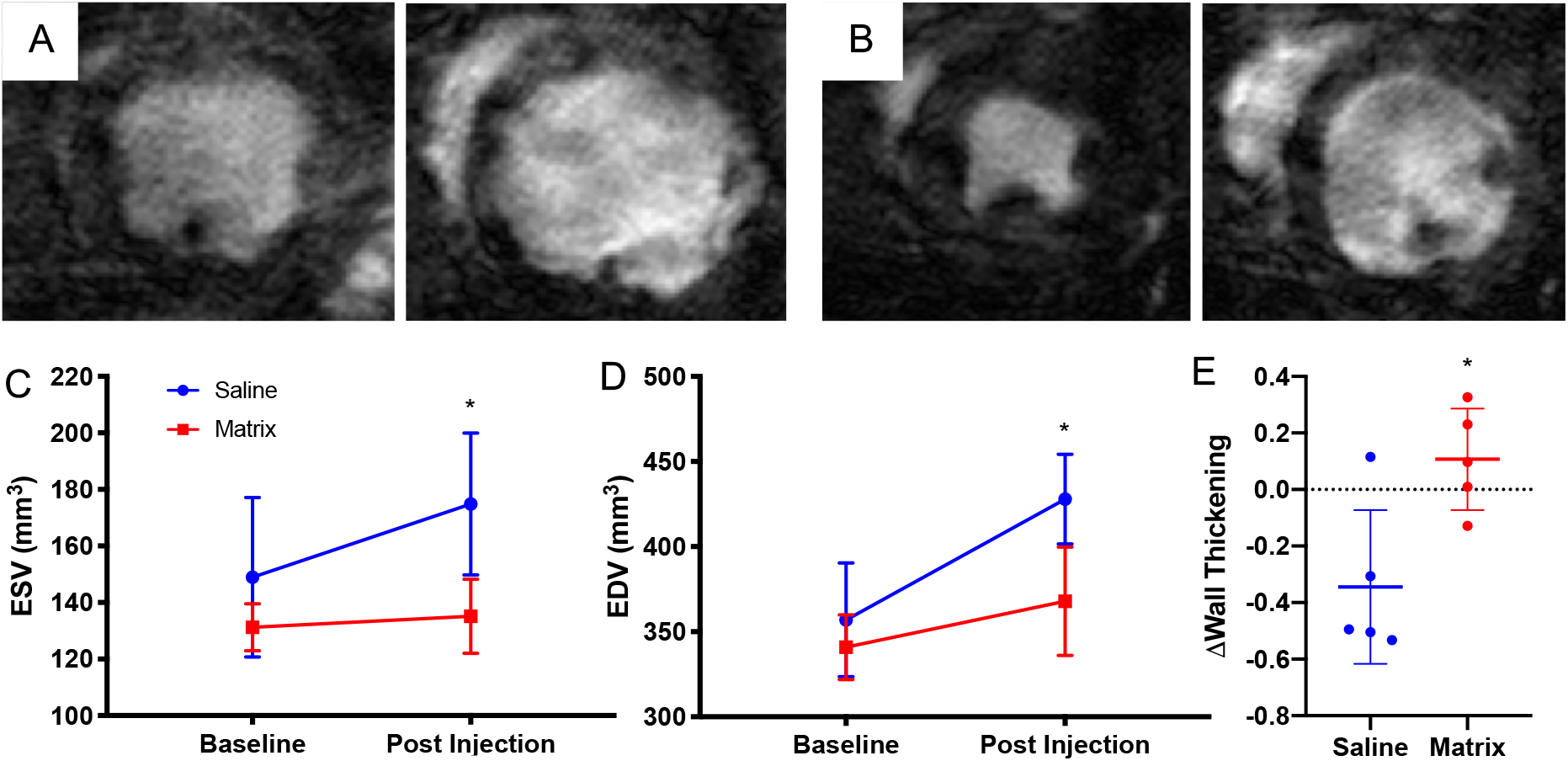
Cardiac Magnetic Resonance Imaging shows the myocardial matrix hydrogel preserves LV volumes and improves cardiac wall thickening up to 4 weeks post-injection. Representative MR images of saline control (A) and matrix treated (B) animals at 4 weeks post-injection are shown at end systole (left) and end diastole (right). MR images were used to determine changes in (C) LVESV and (D) LVEDV from baseline (2 days pre-injection, 8 weeks post-MI) and 4 weeks post-injection (12 weeks post-MI). (E) Change in wall thickening from baseline to 4 weeks post-injection (Δwall thickening) shows the matrix group was significantly greater than saline controls. *p<0.05 compared to baseline values in (C/D) and compared to saline in (E).

**Figure 3:**
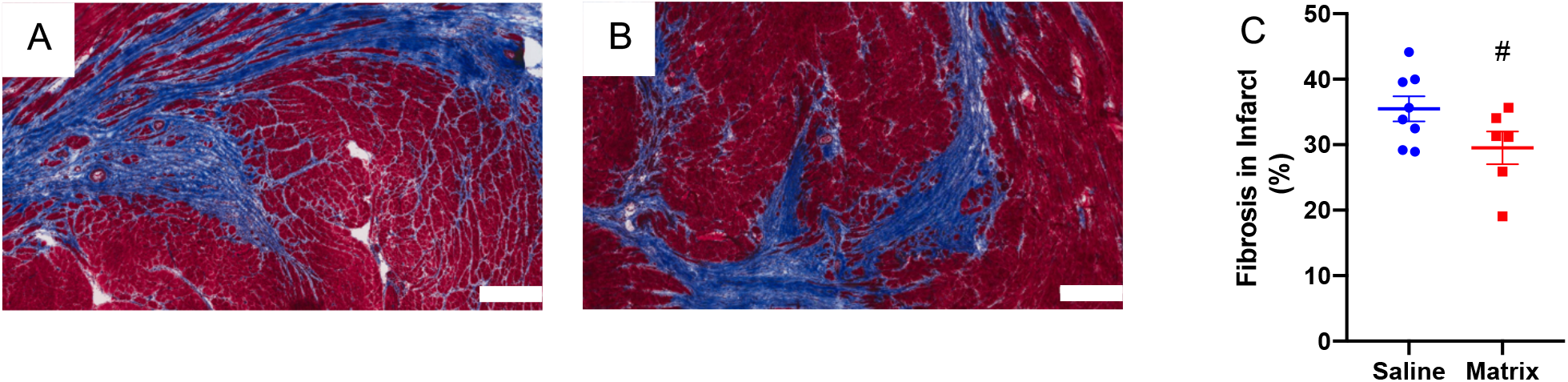
Myocardial matrix modulates the fibrotic response in the infarct. Representative Masson’s Trichrome staining of saline injected (A) and matrix injected (B) infarcts (scale bar =300 μm). Quantification of infarct fibrosis (C) (#p=0.07).

### Effects of Myocardial Matrix on the Cardiac Fibroblasts In Vitro

Cardiac fibroblasts are a dominant cell type in the heart and play a vital role in fibrosis and ventricular remodeling post-MI and throughout HF (21–24). Since the observed reduction in infarct fibrosis (Figure 3) could be a result of the hydrogel modulating fibroblast activity, we evaluated *in vitro* how cardiac fibroblasts interacted with the different forms of the matrix that they would be exposed to *in vivo:* full hydrogel, elution fraction, and digestion fraction (Figure 4a). The elution fraction is the soluble fraction burst released after hydrogel gelation and the digestion fraction is the soluble fraction released after MMP degradation, which would occur upon cell infiltration *in vivo.* MMP-2 and MMP-9 were chosen for this process since they are secreted by both fibroblasts and immune cells, which are known to infiltrate the material (6, 25). Media from each culture condition were then analyzed by gel zymography, which allows for quantification of pro and cleaved MMP-2 and MMP-9. Pro-MMPs must be cleaved in order for their enzymatic function to be activated. Compared to collagen hydrogels, which were used as a control to mimic the infarct scar, cardiac fibroblasts secreted significantly more MMP-2 and MMP-9 when cultured on myocardial matrix hydrogels (Figure 4b, c). Similarly, concentration of cleaved MMP-2 was significantly higher in media collected from fibroblasts exposed to both the elution and digestion fractions compared to their respective enzyme controls (Figure 4d, e).

**Figure 4:**
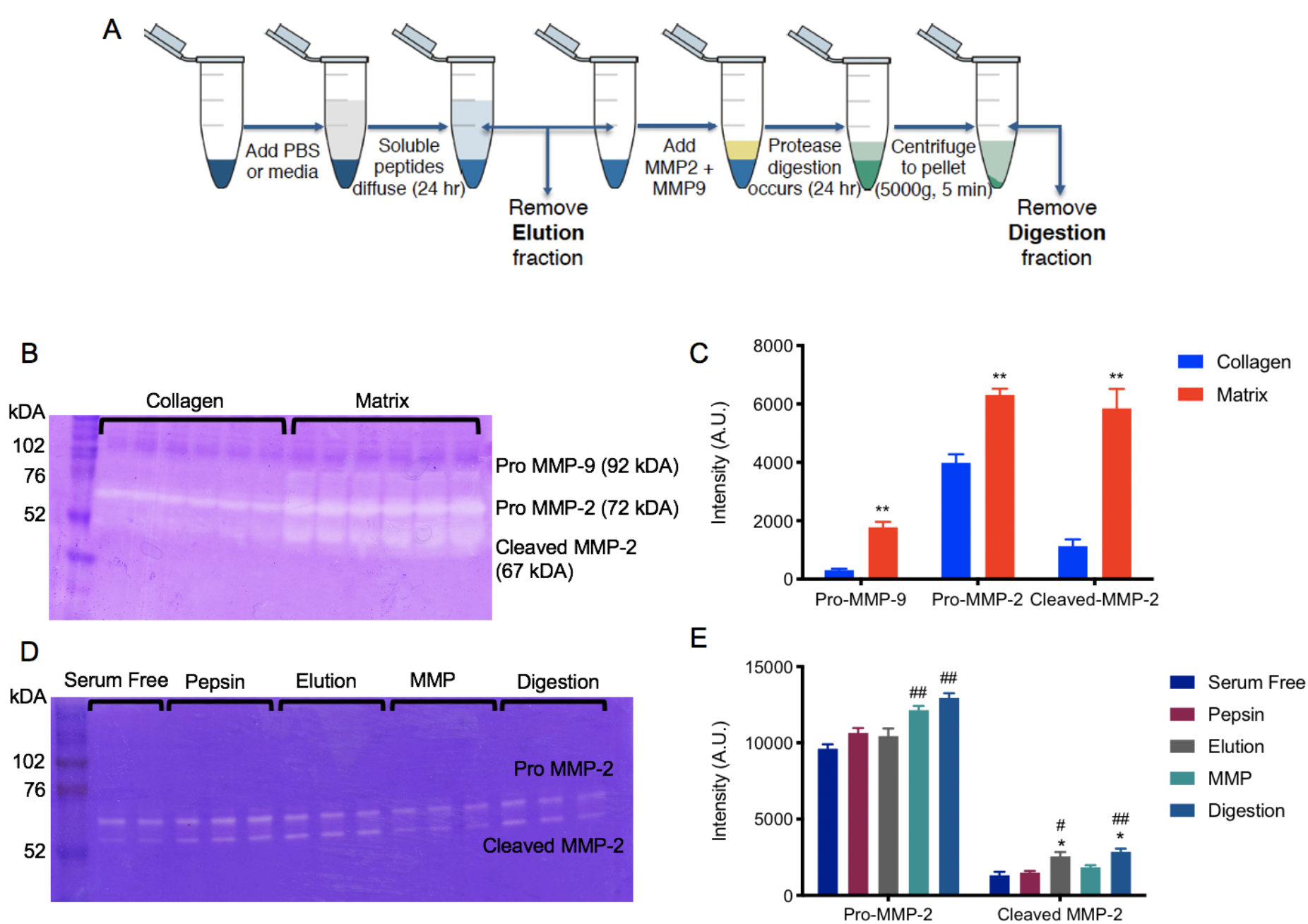
Myocardial matrix induces MMP secretion by cardiac fibroblasts *in vitro.* (A) Schematic depicting simulation of myocardial matrix hydrogel degradation *in vitro.* The myocardial matrix hydrogel is formed, depicted in dark blue. The elution fraction is defined as the soluble peptides that diffuse from the matrix and is illustrated in light blue. After the elution fraction is removed, MMP-2 and MMP-9 are added to matrix. The digestion fraction represents peptides released by the MMP digestion, shown in light green. (B) Zymogram of media collected from cardiac fibroblasts cultured on either collagen or myocardial matrix hydrogels. (C) Analysis of band intensity. (D) Zymogram of media collected from cardiac fibroblasts cultured on tissue culture plastic, with serum free media or post-gelation fractions at a 1:10 dilution. (E) Analysis of band intensity. * p < 0.05, ** p < 0.01; *: compared to collagen in (C) or appropriate enzyme only control in (E), #: compared to serum free media.

### Gene Expression Analysis

A custom NanoString nCounter codeset of 380 genes was designed to provide insight into the mechanism of action of the myocardial matrix hydrogel in a model of chronic MI. The panel was designed with the following pathways in mind: fibrosis, immune response (cytokine profile, transcriptional regulation of macrophages and T-cells), cardiac muscle contraction and development, angiogenesis, apoptosis, and cardiac metabolism using literature to determine key genes from each pathway (6, 22, 26–34). The distribution of genes based on these pathways of interest as well as a full gene list can be seen in Supplementary Tables 3 and 4. Differential gene expression analyses were performed at 1 week post-injection, which is a time when there is peak cell infiltration into the material and when there were significant shifts in global gene expression in a sub-acute MI model (6). After 1 week post-injection of either matrix or saline, we saw clustering based on fold change of gene expression of 41 significantly differentially expressed genes (Figure 5). Evaluating these differentially expressed genes based on a Kyoto Encyclopedia of Genes and Genomes (KEGG) pathway enrichment analysis showed hits associated with HF relevant pathways such as dilated cardiomyopathy, hypertrophic cardiomyopathy, cardiac muscle contraction, adrenergic signaling in cardiomyocytes, and TGF-beta signaling (Table 1). Further details, as well as the full list of KEGG pathways and complete list of genes on the nCounter panel can be found in the Supplemental Appendix (Table S4). At 1 week post-injection, there was a significant increase in the expression of ATPase sarcoplasmic/endoplasmic reticulum Ca2+ transporting 1 (Atp2a1), which is crucial for cardiac muscle contraction (Table 1). We also saw significant downregulation of transforming growth factor-β (Tgfb3), bone morphogenetic protein-2 (Bmp2), and Bmp4 (Figure 5). These genes are involved in the master regulation of fibrogenesis in HF via TGF-β signaling pathways (Table 1) (29, 35). Additionally, we showed downregulation of matrix metalloproteinase 2 (Mmp2) and tissue inhibitor of Mmp2 (Timp2), both of which are associated with fibrogenesis in HF (Figure 5) (25, 36), as well as downregulation of matrix metalloproteinase 2 (Mmp2) and tissue inhibitor of Mmp2 (Timp2), both of which are associated with fibrogenesis in HF (Figure 5). We also observed downregulation of Tlr2, Fos, Akt3, Fcer1a, CD68, Cxcl11, and Pf4 and upregulation of Ccl11 showing modulation of the immune response towards an anti-inflammatory response (Figure 5) (32). Finally, we saw downregulation of key metabolic genes such as Pdk3 and Akt3, which are involved in glucose oxidation.

**Figure 5.**
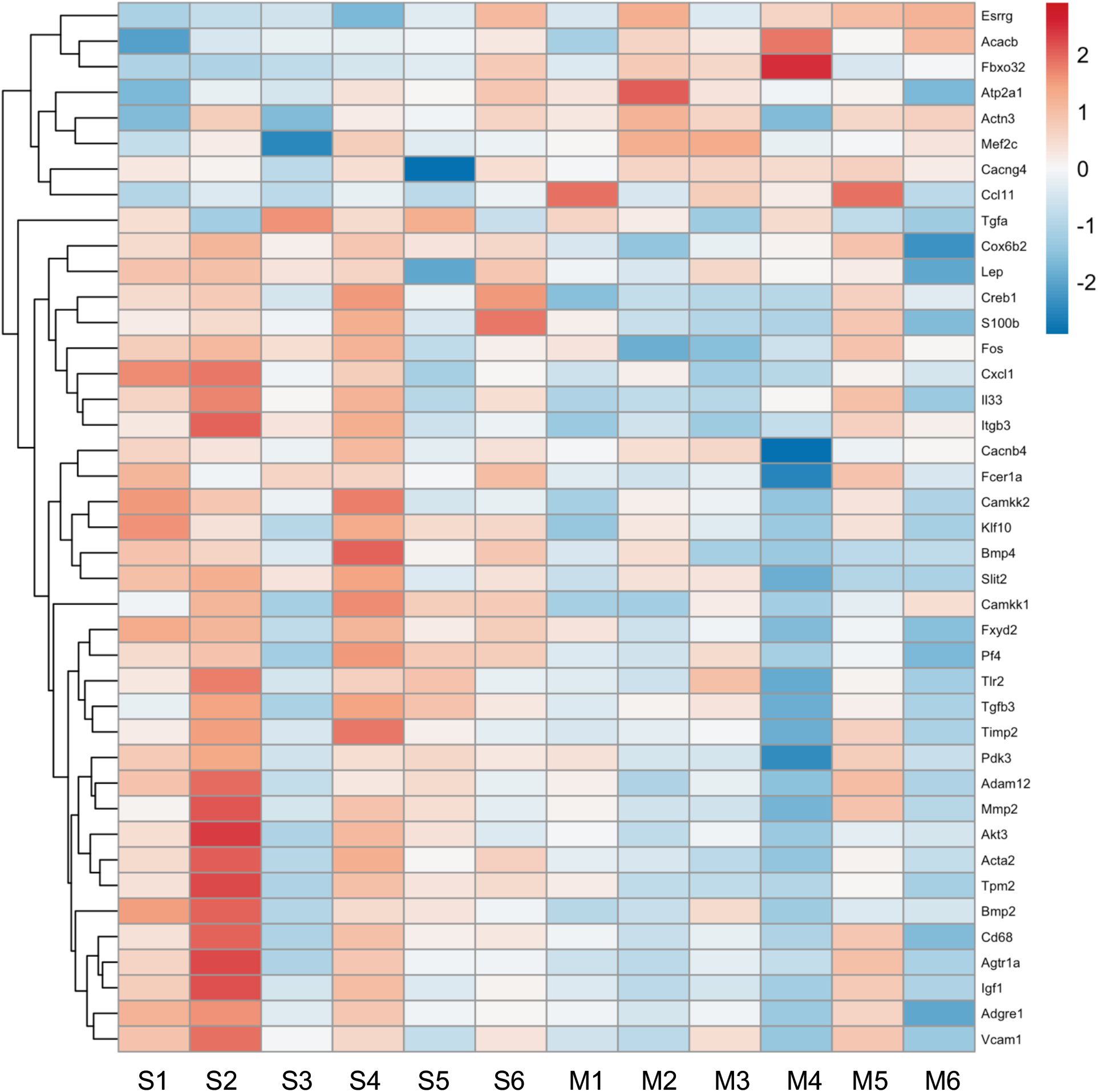
Differential gene expression seen at 1 week post-injection of matrix. Heatmap of 42 significantly differentially expressed genes show that gene expression in the matrix treatment group (M1-6) is distinct from the saline control group (S1-6) at 1 week post-injection.

**Table 1.** KEGG pathway targets and the associated differentially expressed genes.

| Pathway ID | Description | p.adjust | geneID |
| --- | --- | --- | --- |
| rno05410 | Hypertrophic cardiomyopathy (HCM) | 3.5E-06 | Atp2a1/Cacnb4/Cacng4/Igf1/Itgb3/Tgfb3/Tpm2 |
| rno05414 | Dilated cardiomyopathy (DCM) | 3.5E-06 | Atp2a1/Cacnb4/Cacng4/Igf1/Itgb3/Tgfb3/Tpm2 |
| rno04261 | Adrenergic signaling in cardiomyocytes | 3.5E-06 | Agtr1a/Akt3/Atp2a1/Cacnb4/Cacng4/Creb1/Fxyd2/Tpm2 |
| rno04260 | Cardiac muscle contraction | 3.44E-05 | Atp2a1/Cacnb4/Cacng4/Cox6b2/Fxyd2/Tpm2 |
| rno05418 | Fluid shear stress and atherosclerosis | 5.94E-05 | Akt3/Bmp4/Fos/Itgb3/Mef2c/Mmp2/Vcam1 |
| rno04060 | Cytokine-cytokine receptor interaction | 0.000163 | Bmp2/Bmp4/Ccl11/Cxcl1/Il33/Lep/Pf4/Tgfb3 |
| rno04152 | AMPK signaling pathway | 0.000175 | Acacb/Akt3/Camkk2/Creb1/Igf1/Lep |
| rno05412 | Arrhythmogenic right ventricular cardiomyopathy (ARVC) | 0.000175 | Actn3/Atp2a1/Cacnb4/Cacng4/Itgb3 |
| rno04010 | MAPK signaling pathway | 0.000209 | Akt3/Cacnb4/Cacng4/Fos/Igf1/Mef2c/Tgfa/Tgfb3 |
| rno04022 | cGMP-PKG signaling pathway | 0.000594 | Agtr1a/Akt3/Atp2a1/Creb1/Fxyd2/Mef2c |
| rno04668 | TNF signaling pathway | 0.000845 | Akt3/Creb1/Cxcl1/Fos/Vcam1 |
| rno04657 | IL-17 signaling pathway | 0.003297 | Ccl11/Cxcl1/Fos |
| rno04024 | cAMP signaling pathway | 0.007044 | Akt3/Atp2a1/Creb1/Fos/Fxyd2 |
| rno04151 | PI3K-Akt signaling pathway | 0.009474 | Akt3/Creb1/Igf1/Itgb3/Tgfa/Tlr2 |
| rno04062 | Chemokine signaling pathway | 0.01877 | Akt3/Ccl11/Cxcl1/Pf4 |
| rno04620 | Toll-like receptor signaling pathway | 0.022124 | Akt3/Fos/Tlr2 |
| rno04350 | TGF-beta signaling pathway | 0.022252 | Bmp2/Bmp4/Tgfb3 |
| rno04922 | Glucagon signaling pathway | 0.025868 | Acacb/Akt3/Creb1 |
| rno04960 | Aldosterone-regulated sodium reabsorption | 0.03179 | Fxyd2/Igf1 |
| rno04973 | Carbohydrate digestion and absorption | 0.043331 | Akt3/Fxyd2 |

## Discussion

Previous pre-clinical studies have shown efficacy of the myocardial matrix hydrogel in preventing negative LV remodeling and promoting repair in the infarct region in small and large animal subacute MI models in small and large animals (4, 6, 20, 37). This work led to completion of a Phase I clinical trial that also suggested the hydrogel may have potential efficacy in chronic MI and ischemic HF patients (7). This current study therefore aimed to show the efficacy and mechanism by which the hydrogel can repair a chronic MI using a rat model. When the matrix was injected 8 weeks post-MI in the rat model, we observed mitigation of further negative LV remodeling and reduced fibrosis (Figure 2,3). Based on these *in vivo* results, we next turned to *in vitro* assays to detect MMP production by cardiac fibroblasts in the presence of the matrix and its degradation products. These data suggested the matrix may be able to stimulate remodeling of the infarct scar (Figure 4). Finally, *in vivo* gene expression analyses in the chronic MI model showed the importance of differential expression of pathways dysregulated in HF and suppression of chronic inflammation (Figure 5, Table 1).

To first confirm efficacy of the matrix in a chronic MI model *in vivo,* we used MRI to determine changes in cardiac function prior to injection and at 4 weeks post-injection. We demonstrated preserved LV volumes and improved apical wall thickening at 4 weeks post-injection in matrix injected animals (Figure 2). These results support what has been shown in the Phase I Clinical Trial in which the preliminary functional data of LV volumes by MRI analysis suggested functional efficacy in late (chronic) MI patients (7). Histological assessments showed a trend for reduced fibrosis in the infarct of matrix treated animals (Figure 3), suggesting a possible mechanism of action for preserved cardiac function.

Cardiac fibroblasts are one of the dominant cell types of the heart, playing an important role in regulating the mechanics and structure of the myocardium through production of ECM (21, 23). They are also the main cell type responsible for cardiac fibrosis and scar formation in the infarct post MI. Cardiac fibroblasts are critical to the pathogenesis of HF since they are particularly responsive to inflammatory cytokines post-MI and, in turn, produce a wide range of ECM proteins, MMPs, and other paracrine signaling for months to years after MI (21–24). Moreover, the chronic scar is a dynamic tissue in which there are fibrogenic signals and collagen is turned over (38). To better understand the effects of the matrix on cardiac fibroblast activation towards a pro-remodeling vs. fibrotic phenotype, we evaluated how cardiac fibroblasts interacted with the different forms of the matrix *in vitro.* Here we saw significantly more MMP-2/9 secretion by cardiac fibroblasts in response to the matrix and its degradation products, in addition to increased activity of MMP-2. While activation of MMPs acutely post-MI has been associated with progression of LV remodeling due to degradation of native cardiac ECM and subsequent deposition of fibrotic proteins, we have not observed an increase in fibrosis in previous applications of the matrix in sub-acute MI, nor in the chronic MI model presented here (Figure 4) (6, 20, 39, 40). Instead, matrix induced activation of cardiac fibroblasts to secrete MMPs may play a role in reducing fibrosis by actively remodeling the infarct scar.

The NanoString custom codeset was designed as a screening tool to be used to provide an initial understanding of what the mechanism the matrix may have in a chronic MI model, on a gene expression level. Specifically, we saw upregulation of key genes involved in the cardiac muscle contraction pathway such as Atp2a1, otherwise known as SERCA1 (Figure 5). SERCA1 has previously been shown to improve cardiac muscle contraction in small animal models and other isoforms of SERCA genes have been focused on as a targets for gene therapy in HF (41–43). In sub-acute MI it has been shown that the matrix reduces fibrosis up to 5 weeks post-injection, however the mechanism was not understood (6). In this study we observed downregulation of three key genes involved in TGF-β signaling (Tgfb3, Bmp2, Bmp4). TGF-β is a master modulator of fibrogenesis in various diseases and expression of genes in this pathway are positively correlated with fibrosis (29, 35). The matrix may mitigate fibrogenesis through the suppression of TGF-β signaling.

Previous studies on the effect of the matrix on cardiac metabolism have shown a rescue of genes involved in oxidative phosphorylation (6). This is beneficial as cardiomyocyte metabolism is notoriously dysregulated in HF with the suppression of oxidative phosphorylation and enhanced dependence on glucose oxidation for ATP supply contributing to adverse remodeling. In this study of chronic MI we did not see upregulation of genes that regulate oxidative phosphorylation directly, but we found downregulation of glycolysis/glucose oxidation genes such as Pdk3 and Akt3. These genes are regulators of glucose oxidation and their chronic upregulation in HF is linked to negative LV remodeling (44–46). Suppression of these genes suggest the matrix is able to modulate cardiac metabolism in chronic MI.

In sub-acute MI, there is evidence that the matrix is able to modulate the sub-acute inflammatory and immune response to promote injury repair, reduce apoptosis of at-risk cardiomyocytes, and promote neovascularization (27, 47, 48). Interestingly, in this study we demonstrated an alternate modulation of the inflammatory and immune response. In matrix-treated animals there was reduced expression of Mmp2 and Timp2, which are both associated with chronic inflammation in HF leading to fibrosis, increased wall stiffness, and reduced contractility (36). While the *in vitro* data suggests the matrix is able to promote MMP-2 production (Figure 4), this assay only assessed cardiac fibroblast MMP production and the *in vivo* gene expression analysis takes into account all cell types located in the infarct region of the myocardium such as immune cells, which are known to produce MMPs (28). The gene expression analysis shows significant downregulation of inflammatory cytokines such as Tlr2, Fos, Akt3, Cxcl1 and Pf4 as well as Cd68 and Fcer1a, suggesting reduced migration of immune cells in matrix treated animals. A KEGG pathway enrichment analysis shows these genes are involved in key immune response pathways such as: cytokine-cytokine interaction, TNF signaling, toll like receptor signaling, and IL-17 signaling, all of which are known to contribute to adverse remodeling throughout HF (Table 1) (27, 32, 49). This suggests the *in vivo* reduction in Mmp2/Timp2 may be due to the suppression of inflammation and the pro-remodeling immune response induced by the matrix. While acute inflammation and immune response immediately after MI is required to some extent for injury repair, chronic inflammation in HF is associated with the negative feedback loop of continued adverse remodeling (32, 49, 50). Overall, this study provides evidence that the matrix is able to modulate pathways of cardiac muscle contraction, fibrosis, metabolism, and the inflammatory/immune response in a small animal model of chronic MI to preserve cardiac function and prevent progression of adverse LV remodeling.

### Study Limitations

This study is limited by the use of a small animal murine model of chronic MI. While ischemia reperfusion is more representative of the patient population, which is revascularized, the model is not as severe as a permanent ligation model and therefore, this may have resulted in smaller differences in function and gene expression. Additionally, the nCounter custom codeset was designed for this project with genes across many different pathways for screening for potential mechanisms of action, which limits the analysis in delineating in-depth details of pathway modulation from the material in a chronic MI environment.

### Conclusion

This study is the first investigation into the efficacy and mechanism of a myocardial matrix hydrogel to prevent worsening HF in a small animal model of chronic MI. We observed preserved LV volumes and apical wall thickening up to 4 weeks post injection of the hydrogel, as well as trends for reduced fibrosis. Further, we found the hydrogel modulated key pathways dysregulated in HF such as dilated cardiomyopathy, hypertrophic cardiomyopathy, cardiac muscle contraction, adrenergic signaling in cardiomyocytes, TGF-β signaling, and inflammation.

## Supporting information

Supplemental Index

## Acknowledgments

The authors would like to thank Pamela Duran for feedback on the manuscript, Dr. Raymond Wang for feedback on the manuscript and help with R script and NanoString data analysis, and Dr. Elsa Molina of the Sanford Consortium Genomics Core for NanoString analysis prep and troubleshooting.

## References

1. Benjamin EJ, Muntner P, Alonso A, Bittencourt MS, Callaway CW, Carson AP, et al. Heart Disease and Stroke Statistics-2019 Update: A Report From the American Heart Association. Circulation. 2019;139:(10):e56–e528.

2. Diaz MD, Christman KL. Injectable Hydrogels to Treat Myocardial Infarction. Cardiovascular Regenerative Medicine 2019. p. 185–206.

3. Venugopal JR, Prabhakaran MP, Mukherjee S, Ravichandran R, Dan K, Ramakrishna S. Biomaterial strategies for alleviation of myocardial infarction. J R Soc Interface. 2012;9(66):1–19.

4. Singelyn JM, Sundaramurthy P, Johnson TD, Schup-Magoffin PJ, Hu DP, Faulk DM, et al. Catheter-deliverable hydrogel derived from decellularized ventricular extracellular matrix increases endogenous cardiomyocytes and preserves cardiac function post-myocardial infarction. J Am Coll Cardiol. 2012;59(8):751–63.

5. Sonya B. Seif-Naraghi JMS, Michael A. Salvatore,2 Kent G. Osborn,, Jean J. Wang US, Oi Ling Kwan, G. Monet Strachan, Jonathan Wong,, Pamela J. Schup-Magoffin RLB, Kendra Bartels, Jessica A. DeQuach,, Mark Preul AMK, Anthony N. DeMaria, Nabil Dib, Karen L. Christman. Safety and Efficacy of an Injectable Extracellular Matrix Hydrogel for Treating Myocardial Infarction. Science Translational Medicine. 2013;5(173).

6. Wassenaar JW, Gaetani R, Garcia JJ, Braden RL, Luo CG, Huang D, et al. Evidence for Mechanisms Underlying the Functional Benefits of a Myocardial Matrix Hydrogel for Post-MI Treatment. J Am Coll Cardiol. 2016;67(9):1074–86.

7. Traverse JH, Henry TD, Dib N, Patel AN, Pepine C, Schaer GL, et al. First-in-Man Study of a Cardiac Extracellular Matrix Hydrogel in Early and Late Myocardial Infarction Patients. JACC Basic Transl Sci. 2019;4(6):659–69.

8. Ungerleider JL, Johnson TD, Rao N, Christman KL. Fabrication and characterization of injectable hydrogels derived from decellularized skeletal and cardiac muscle. Methods. 2015;84:53–9.

9. Hernandez MJ YG, Zelus EI, et al. Manufacturing considerations for producing and assessing decellularized extracellular matrix hydrogels. Methods. 2019(171):20–7.

10. Hill RC, Calle EA, Dzieciatkowska M, Niklason LE, Hansen KC. Quantification of extracellular matrix proteins from a rat lung scaffold to provide a molecular readout for tissue engineering. Molecular and Cellular Proteomics. 2015;14(4):961–73.

11. Singelyn JM, DeQuach JA, Seif-Naraghi SB, Littlefield RB, Schup-Magoffin PJ, Christman KL. Naturally derived myocardial matrix as an injectable scaffold for cardiac tissue engineering. Biomaterials. 2009;30(29):5409–16.

12. Manuel D. Cerqueira MNJW, MD; Vasken Dilsizian, MD; Alice K. Jacobs, MD; Sanjiv Kaul, MD; Warren K. Laskey, MD; Dudley J. Pennell, MD; John A. Rumberger, MD; Thomas Ryan, MD; Mario S. Verani, MD. Standardized Myocardial Segmentation and Nomenclature for Tomographic Imaging of the Heart. Circulation. 2002;105:539.

13. Ulrika S Pahlm JFU, Einar Heiberg, Henrik Engblom, David Erlinge, Matthias Götberg and Håkan Arheden. Regional wall function before and after acute myocardial infarction; an experimental study in pigs. BMC Cardiovascular Disorders 2014;14:10.

14. Yuan Ye K, Sullivan KE, Black LD. Encapsulation of cardiomyocytes in a fibrin hydrogel for cardiac tissue engineering. J Vis Exp. 2011(55).

15. Jardine L, Wiscombe S, Reynolds G, McDonald D, Fuller A, Green K, et al. Lipopolysaccharide inhalation recruits monocytes and dendritic cell subsets to the alveolar airspace. Nature Communications. 2019;10(1):1999.

16. Hussain S, Johnson CG, Sciurba J, Meng X, Stober VP, Liu C, et al. TLR5 participates in the TLR4 receptor complex and promotes MyD88-dependent signaling in environmental lung injury. Elife. 2020;9.

17. Wang H, Horbinski C, Wu H, Liu Y, Sheng S, Liu J, et al. NanoStringDiff: a novel statistical method for differential expression analysis based on NanoString nCounter data. Nucleic Acids Research. 2016;44(20):e151–e.

18. Yu G, Wang L-G, Han Y, He Q-Y. clusterProfiler: an R Package for Comparing Biological Themes Among Gene Clusters. OMICS: A Journal of Integrative Biology. 2012;16(5):284–7.

19. Luo W, Brouwer C. Pathview: an R/Bioconductor package for pathway-based data integration and visualization. Bioinformatics. 2013;29(14):1830–1.

20. Seif-Naraghi SB, Singelyn JM, Salvatore MA, Osborn K, Wang JJ, Dequach JA, et al. Safety and efficacy of an injectable extracellular matrix hydrogel for treating myocardial infarction in pre-clinical animal studies. Science Translational Medicine. 2013;5:173ra25.

21. Zhou P, Pu WT. Recounting Cardiac Cellular Composition. Circ Res. 2016;118(3):368–70.

22. Nagaraju CK, Robinson EL, Abdesselem M, Trenson S, Dries E, Gilbert G, et al. Myofibroblast Phenotype and Reversibility of Fibrosis in Patients With End-Stage Heart Failure. J Am Coll Cardiol. 2019;73(18):2267–82.

23. Porter KE, Turner NA. Cardiac fibroblasts: At the heart of myocardial remodeling. Pharmacol Ther. 2009;123(2):255–78.

24. Turner NA, Porter KE. Regulation of myocardial matrix metalloproteinase expression and activity by cardiac fibroblasts. IUBMB Life. 2012;64(2):143–50.

25. DeLeon-Pennell KY, Meschiari CA, Jung M, Lindsey ML. Matrix Metalloproteinases in Myocardial Infarction and Heart Failure. Prog Mol Biol Transl Sci. 2017;147:75–100.

26. Doenst T, Nguyen TD, Abel ED. Cardiac metabolism in heart failure: implications beyond ATP production. Circ Res. 2013;113(6):709–24.

27. Frangogiannis NG. The inflammatory response in myocardial injury, repair, and remodelling. Nat Rev Cardiol. 2014;11(5):255–65.

28. Frangogiannis NG. Pathophysiology of Myocardial Infarction. Compr Physiol. 2015;5(4):1841–75.

29. Dobaczewski M, Chen W, Frangogiannis NG. Transforming growth factor (TGF)-beta signaling in cardiac remodeling. J Mol Cell Cardiol. 2011;51(4):600–6.

30. Khalil H, Kanisicak O, Prasad V, Correll RN, Fu X, Schips T, et al. Fibroblast-specific TGF-beta-Smad2/3 signaling underlies cardiac fibrosis. J Clin Invest. 2017;127(10):3770–83.

31. Noordali H, Loudon BL, Frenneaux MP, Madhani M. Cardiac metabolism - A promising therapeutic target for heart failure. Pharmacol Ther. 2018;182:95–114.

32. Riehle C, Bauersachs J. Key inflammatory mechanisms underlying heart failure. Herz. 2019;44(2):96–106.

33. Gaballa MA, Goldman S. Ventricular remodeling in heart failure. Journal of Cardiac Failure. 2002;8(6):S476–S85.

34. Thomas J Cahill RKK. Heart failure after myocardial infarction in the era of primary percutaneous coronary intervention: Mechanisms, incidence and identification of patients at risk. World Journal of Cardiology. 2017;9(5).

35. Biernacka A, Dobaczewski M, Frangogiannis NG. TGF-beta signaling in fibrosis. Growth Factors. 2011;29(5):196–202.

36. Kobusiak-Prokopowicz M, Krzysztofik J, Kaaz K, Jolda-Mydlowska B, Mysiak A. MMP-2 and TIMP-2 in Patients with Heart Failure and Chronic Kidney Disease. Open Med (Wars). 2018;13:237–46.

37. Sonya B. Seif-Naraghi BSE, Michael A. Salvatore, B.S., Pam J. Schup-Magoffin, B.A.,, Diane P. Hu MS, and Karen L. Christman, Ph.D. Design and Characterization of an Injectable Pericardial Matrix Gel-A Potentially Autologous Scaffold for Cardiac Tissue Engineering. Tissue Eng Part A. 2010;16(6).

38. Yao Sun KTW. Infarct Scar: A dynamic tissue. Cardiovascular Research. 2000;46:6.

39. Singelyn JM, Sundaramurthy P, Johnson TD, Schup-Magoffin PJ, Hu DP, Faulk DM, et al. Catheter-deliverable hydrogel derived from decellularized ventricular extracellular matrix increases endogenous cardiomyocytes and preserves cardiac function post-myocardial infarction. Journal of the American College of Cardiology. 2012;59(8):751–63.

40. Spinale FG. Matrix Metalloproteinases: Regulation and Dysregulation in the Failing Heart. Circulation Research. 2002;90(5):520–30.

41. M. Jane Lalli JY, Vikram Prasad, Katsuji Hashimoto, Dave Plank, Gopal J. Babu, Darryl Kirkpatrick, Richard A. Walsh, Mark Sussman, Atsuko Yatani, Eduardo Marbán, Muthu Periasamy. Sarcoplasmic Reticulum Ca2+ ATPase (SERCA) 1a Structurally Substitutes for SERCA2a in the Cardiac Sarcoplasmic Reticulum and Increases Cardiac Ca2+ Handling Capacity. Circulation Research. 2001;89:7.

42. Teucher N, Prestle J, Seidler T, Currie S, Elliott EB, Reynolds DF, et al. Excessive sarcoplasmic/endoplasmic reticulum Ca2+-ATPase expression causes increased sarcoplasmic reticulum Ca2+ uptake but decreases myocyte shortening. Circulation. 2004;110(23):3553–9.

43. Eisner D, Caldwell J, Trafford A. Sarcoplasmic reticulum Ca-ATPase and heart failure 20 years later. Circ Res. 2013;113(8):958–61.

44. Wende AR, O’Neill BT, Bugger H, Riehle C, Tuinei J, Buchanan J, et al. Enhanced cardiac Akt/protein kinase B signaling contributes to pathological cardiac hypertrophy in part by impairing mitochondrial function via transcriptional repression of mitochondrion-targeted nuclear genes. Mol Cell Biol. 2015;35(5):831–46.

45. Taniyama Y, Ito, M., Sato, K., Kuester, C., Veit, K., Tremp, G., Liao, R., Colucci, W. S., Ivashchenko, Y., Walsh, K., & Shiojima, I. Akt3 overexpression in the heart results in progression from adaptive to maladaptive hypertrophy. Journal of molecular and cellular cardiology. 2005;38(2):10.

46. Gray LR, Tompkins SC, Taylor EB. Regulation of pyruvate metabolism and human disease. Cell Mol Life Sci. 2014;71(14):2577–604.

47. Porur Somasundaram GR, Himanshu Nagar, Daniela Kraemer, Leonardo Mendoza, Lloyd H Michael, George H Caughey, Mark L Entman, and Nikolaos G Frangogiannis. Mast cell tryptase may modulate endothelial cell phenotype in healing myocardial infarcts. Journal of Pathology. 2008;205:9.

48. Bischoff SC. Role of mast cells in allergic and non-allergic immune responses: comparison of human and murine data. Nat Rev Immunol. 2007;7(2):93–104.

49. Ong SB, Hernandez-Resendiz S, Crespo-Avilan GE, Mukhametshina RT, Kwek XY, Cabrera-Fuentes HA, et al. Inflammation following acute myocardial infarction: Multiple players, dynamic roles, and novel therapeutic opportunities. Pharmacol Ther. 2018;186:73–87.

50. Aurora AB, Porrello ER, Tan W, Mahmoud AI, Hill JA, Bassel-Duby R, et al. Macrophages are required for neonatal heart regeneration. J Clin Invest. 2014;124(3):1382–92.

