## Supplemental Index for "Injectable Myocardial Matrix Hydrogel Mitigates Negative Left Ventricular Remodeling in a Chronic Myocardial Infarction Model"

**Supplemental Appendix:**

**Table S1.** Abundance of ECM protein components determined from targeted Quantitative conCATamers and Liquid Chromatography - Selected Reaction Monitoring (LC-SRM) analysis.

| **Protein** | **Gene** | **Functional Classification** | **Protein Abundance [nmol/g]** |
| --- | --- | --- | --- |
| Collagen alpha-1(I) chain | COL1A1 | Fibrillar Collagen | 719.69 |
| Collagen alpha-2(I) chain | COL1A2 | Fibrillar Collagen | 398.54 |
| Collagen alpha-1(I) chain(fragment) | COL1A1 | Fibrillar Collagen | 193.54 |
| Collagen alpha-3(VI) chain | COL6A3 | Matricellular | 125.10 |
| Collagen alpha-1(VI) chain | COL6A1 | Matricellular | 74.36 |
| Collagen alpha-1/5(IV) chain(Arresten/Core Protein) | COL4A1/5 | Basement Membrane | 62.51 |
| Collagen alpha-1(V) chain | COL5A1 | Fibrillar Collagen | 29.55 |
| Collagen alpha-1(IV) chain(Arresten/Core Protein) | COL4A1 | Basement Membrane | 21.85 |
| Perlecan | HSPG2 | Basement Membrane | 21.73 |
| fibronectin 1(type-III 4 domain) | FN1 | Matricellular | 20.65 |
| fibronectin 1(type-III 7 domain) | FN1 | Matricellular | 20.47 |
| Collagen alpha-2(IV) chain(Canstatin/Core Protein) | COL4A2 | Basement Membrane | 12.70 |
| Collagen alpha-2(V) chain | COL5A2 | Fibrillar Collagen | 9.36 |
| Collagen alpha-2(VI) chain | COL6A2 | Matricellular | 8.59 |
| Fibrillin 1 | FBN1 | Structural ECM | 5.67 |
| Periostin | POSTN | Matricellular | 2.70 |
| Lumican | LUM | Matricellular | 1.91 |
| TnxB Protein | TNXB | Matricellular | 1.67 |
| Fibrillin 2 | FBN2 | Structural ECM | 1.09 |
| Dermatopontin | DPT | Matricellular | 0.91 |
| Laminin Beta-2 | LAMB2 | Basement Membrane | 0.51 |
| Laminin Gamma-1 | LAMC1 | Basement Membrane | 0.47 |
| Laminin Gamma-1 | LAMC1 | Basement Membrane | 0.47 |
| Laminin Beta-2 | LAMB2 | Basement Membrane | 0.45 |
| Emilin 1 | EMILIN1 | Matricellular | 0.41 |
| Collagen alpha-1(XVIII) chain | COL18A1 | Matricellular | 0.17 |
| Fibulin 5 | FBLN5 | Matricellular | 0.14 |
| Laminin alpha-2 | LAMA2 | Basement Membrane | 0.07 |
| Actin (All Isoforms) | ACT | Cytoskeletal | 2.93 |
| Glyceraldehyde-3-phosphate dehydrogenase | GAPDH | Other Cellular | 1.77 |
| Myosin(Myosin-3,4,6,7) | MYH | Cytoskeletal | 0.55 |

**Table S2.** Top 20 Proteins in Decellularized Myocardial Matrix by Global Proteomics

| **Proteins** | **Gene** | **Functional Classification** | **Unique PSM's** |
| --- | --- | --- | --- |
| Perlecan | HSPG2 | Basement Membrane | 176 |
| Laminin subunit gamma-1 | LAMC1 | Basement Membrane | 39 |
| Laminin subunit beta-2 | LAMB2 | Basement Membrane | 48 |
| Laminin alpha-5 | LAMA5 | Basement Membrane | 45 |
| Collagen alpha-1(IV) chain | COL4A1 | Basement Membrane | 37 |
| Collagen alpha-2(I) chain | COL1A2 | Fibrillar Collagen | 221 |
| Collagen alpha-1(III) chain | COL3A1 | Fibrillar Collagen | 220 |
| Collagen alpha-2(V) chain | COL5A2 | Fibrillar Collagen | 69 |
| Collagen alpha-1(I) chain | COL1A1 | Fibrillar Collagen | 228 |
| Collagen alpha-1(V) chain | COL5A1 | Fibrillar Collagen | 38 |
| Collagen alpha-3(VI) chain | COL6A3 | Matricellular | 113 |
| Collagen alpha-2(IV) chain | COL4A2 | Matricellular | 71 |
| TnxB Protein | TNXB | Matricellular | 57 |
| Fibronectin | FN1 | Matricellular | 64 |
| Fibrillin-1 | FBN1 | Structural ECM | 146 |
| Fibrillin-2 | FBN2 | Structural ECM | 39 |

**Table S3.** Design Approach for NanoString nCounter CodeSet

| **Pathway** | **Number of Genes** |
| --- | --- |
| Metabolism | 60 |
| Apoptosis | 23 |
| Neovascularization | 65 |
| Neurogenesis | 24 |
| Immune/Inflammatory Response | 54 |
| Fibrosis | 36 |
| Myogenesis/muscle contraction | 69 |
| Muscle hypertrophy | 44 |
| Housekeeping | 6 |

**Table S4. Full Gene List for NanoString nCounter CodeSet**

| **Gene Identifier** | **Accession Number** |
| --- | --- |
| ABCF1 | NM_001109883.2 |
| Abl1 | NM_001100850.1 |
| Acacb | NM_053922.1 |
| Acly | NM_016987.2 |
| Acox2 | NM_145770.1 |
| Acsm5 | NM_001014162.4 |
| Acta2 | NM_031004.2 |
| Actc1 | NM_019183.1 |
| Actn2 | NM_001170325.1 |
| Actn3 | NM_133424.2 |
| Acvrl1 | NM_022441.2 |
| Adam12 | XM_017590240.1 |
| Adam17 | NM_020306.1 |
| Adgre1 | NM_001007557.1 |
| Adora1 | NM_017155.2 |
| Adrb2 | NM_012492.2 |
| Aggf1 | XM_001060407.6 |
| Agrn | NM_175754.1 |
| Agt | NM_134432.2 |
| Agtr1a | NM_030985.4 |
| Akt1 | NM_033230.1 |
| Akt2 | NM_017093.1 |
| Akt3 | NM_031575.1 |
| Amot | XM_001056974.3 |
| Ang2 | NM_001012359.1 |
| Angptl1 | NM_001109383.1 |
| Angptl3 | NM_001025065.1 |
| Angptl4 | NM_199115.2 |
| Arg1 | NM_017134.2 |
| Artn | NM_053397.1 |
| Atf3 | NM_012912.1 |
| Atf6 | NM_001107196.1 |
| Atp12a | NM_133517.1 |
| Atp1a4 | NM_022848.1 |
| Atp1b1 | NM_013113.2 |
| Atp1b2 | NM_012507.3 |
| Atp1b3 | NM_012913.1 |
| Atp1b4 | NM_053381.1 |
| Atp2a1 | NM_058213.1 |
| Atp2a2 | NM_001110139.2 |
| ATP5F1A | NM_023093.1 |
| Bcl2 | NM_016993.1 |
| Bdh2 | NM_001106473.1 |
| Bmp2 | NM_017178.1 |
| Bmp4 | NM_012827.2 |
| Bmp7 | NM_001191856.1 |
| Bst1 | NM_030848.1 |
| Btg1 | NM_017258.1 |
| Btk | NM_001007798.1 |
| Cacna1c | NM_012517.2 |
| Cacna1d | NM_017298.1 |
| Cacna2d2 | NM_175592.2 |
| Cacnb2 | NM_053851.1 |
| Cacnb3 | NM_012828.2 |
| Cacnb4 | NM_001105733.1 |
| Cacng1 | NM_019255.1 |
| Cacng4 | NM_080692.1 |
| Cacng5 | NM_080693.1 |
| Calm1 | NM_031969.2 |
| Calm2 | NM_017326.2 |
| Camk2d | NM_012519.2 |
| Camkk1 | NM_031662.1 |
| Camkk2 | NM_031338.1 |
| Casp1 | NM_012762.2 |
| Casp3 | NM_012922.2 |
| Casp4 | NM_053736.2 |
| Casp6 | NM_031775.2 |
| Casp7 | NM_022260.3 |
| Casq2 | NM_017131.2 |
| Cat | NM_012520.1 |
| Ccl1 | NM_001191092.1 |
| Ccl11 | NM_019205.1 |
| Ccl2 | NM_031530.1 |
| Ccl3 | NM_013025.2 |
| Ccl9 | NM_001012357.1 |
| Ccn2 | NM_022266.2 |
| Ccr1 | NM_020542.2 |
| Ccr2 | NM_021866.1 |
| Ccr3 | NM_053958.1 |
| Ccr5 | NM_053960.3 |
| Cd36 | NM_031561.2 |
| Cd40 | NM_134360.1 |
| Cd59 | NM_012925.1 |
| Cd68 | NM_001031638.1 |
| Cdc42 | NM_171994.4 |
| Cflar | NM_057138.2 |
| Chat | NM_001170593.1 |
| Chrm2 | NM_031016.1 |
| Cox4i1 | NM_017202.1 |
| Cox4i2 | NM_053472.1 |
| Cox5b | NM_053586.1 |
| Cox6a1 | NM_012814.1 |
| Cox6a2 | NM_012812.3 |
| Cox6b1 | NM_001145273.1 |
| Cox6b2 | NM_001039085.1 |
| Cox6c | NM_019360.2 |
| Cox7a2l | NM_001106704.1 |
| Cox7b2 | XM_006221734.3 |
| Cpt1a | NM_031559.2 |
| Cpt1b | NM_013200.1 |
| Creb1 | NM_134443.1 |
| Crhr2 | NM_022714.1 |
| Crtc2 | NM_001033895.1 |
| Cs | NM_130755.1 |
| Csf3 | NM_017104.1 |
| Ctf1 | NM_017129.1 |
| Cxcl1 | NM_030845.1 |
| Cxcl10 | NM_139089.1 |
| Cxcl2 | NM_053647.1 |
| Cxcl3 | NM_138522.1 |
| Cxcl6 | NM_022214.1 |
| Cxcr4 | NM_022205.3 |
| Dffa | NM_053679.2 |
| Dffb | NM_053362.1 |
| Diablo | NM_001008292.1 |
| Dusp5 | NM_133578.1 |
| Dyrk1a | NM_012791.2 |
| Egf | NM_012842.1 |
| Egfl7 | NM_139104.1 |
| Eif2b5 | NM_138866.2 |
| Ep300 | XM_001076610.3 |
| Erbb2 | NM_017003.2 |
| Ereg | NM_021689.1 |
| Esr1 | NM_012689.1 |
| Esrrg | NM_203336.2 |
| Fabp3 | NM_024162.1 |
| Fadd | NM_152937.2 |
| Fas | NM_139194.2 |
| Faslg | NM_012908.1 |
| Fasn | NM_017332.1 |
| Fbp2 | NM_053716.1 |
| Fbxo32 | NM_133521.1 |
| Fcer1a | NM_012724.4 |
| Fgf2 | NM_019305.2 |
| Fh | NM_017005.2 |
| Flt4 | NM_053652.1 |
| Fmn1 | XM_006224644.3 |
| Fos | NM_022197.1 |
| Foxo1 | NM_001191846.2 |
| Foxo3 | NM_001106395.1 |
| Foxp3 | NM_001108250.1 |
| Fst | NM_012561.1 |
| Fxyd2 | NM_017349.2 |
| G6pd | NM_017006.2 |
| Gadd45a | NM_024127.2 |
| Gata3 | NM_133293.1 |
| Gata4 | NM_144730.1 |
| Gja1 | NM_012567.2 |
| Gls | NM_012569.2 |
| Glud1 | NM_012570.1 |
| Got1 | NM_012571.1 |
| Got2 | NM_013177.2 |
| Gpt | NM_031039.1 |
| Gpt2 | NM_001012057.1 |
| Gpx | NM_030826.2 |
| Gsk3a | NM_017344.1 |
| Gsk3b | NM_032080.1 |
| Gsr | NM_053906.1 |
| Gss | NM_012962.1 |
| GUSB | NM_017015.2 |
| Hey1 | NM_001191845.1 |
| Hey2 | NM_130417.1 |
| Hk2 | NM_012735.1 |
| HPRT1 | NM_012583.2 |
| Hspa5 | NM_013083.1 |
| Idh1 | NM_031510.1 |
| Idh2 | NM_001014161.1 |
| Ifng | NM_138880.2 |
| Igf1 | NM_001082477.2 |
| Il10 | NM_012854.2 |
| Il12a | NM_053390.1 |
| Il13 | NM_053828.1 |
| Il17a | NM_001106897.1 |
| Il1b | NM_031512.1 |
| IL38 | NM_001108571.1 |
| Il1rn | NM_022194.2 |
| Il2 | NM_053836.1 |
| Il23a | NM_130410.2 |
| Il33 | NM_001014166.1 |
| Il36a | NM_001106554.1 |
| Il4 | NM_201270.1 |
| Il5 | NM_021834.1 |
| Il6 | NM_012589.1 |
| Il6st | NM_001008725.3 |
| Irf4 | NM_001106108.1 |
| Irf5 | NM_001106586.1 |
| Itga1 | NM_030994.2 |
| Itga2 | XM_345156.6 |
| Itga3 | XM_003752369.1 |
| Itgav | NM_001106549.1 |
| Itgb1 | NM_017022.2 |
| Itgb3 | NM_153720.1 |
| Itgb5 | NM_147139.2 |
| Itgb6 | NM_001004263.1 |
| Itgb8 | NM_001108726.1 |
| Jdp2 | NM_053894.1 |
| Junb | NM_021836.2 |
| Kcnd2 | NM_031730.2 |
| Kdr | NM_013062.1 |
| Kit | NM_022264.1 |
| Klf10 | NM_031135.2 |
| Kmt2a | XM_006226455.1 |
| LDHA | NM_017025.1 |
| Lep | NM_013076.3 |
| Letm1 | NM_001005884.1 |
| Lifr | NM_031048.1 |
| Lipe | NM_012859.1 |
| Lox | NM_017061.2 |
| Lrg1 | NM_001009717.1 |
| Map2k3 | XM_001077724.1 |
| Mapk1 | NM_053842.1 |
| Mapk11 | NM_001109532.2 |
| Mapk12 | NM_021746.1 |
| Mapk14 | NM_031020.2 |
| Mapk8 | XM_001056513.1 |
| Mapkapk2 | NM_178102.2 |
| Mcl1 | NM_021846.2 |
| Mcu | NM_001106398.1 |
| Mdh1 | NM_033235.1 |
| Mdk | NM_030859.2 |
| Mef2c | XM_006223957.2 |
| Mef2d | NM_030860.2 |
| Mmp1 | NM_001134530.1 |
| Mmp10 | NM_133514.1 |
| Mmp12 | NM_053963.1 |
| Mmp13 | NM_133530.1 |
| Mmp14 | NM_031056.1 |
| Mmp19 | NM_001107159.1 |
| Mmp2 | NM_031054.2 |
| Mmp25 | XM_001055465.2 |
| Mmp28 | NM_001079888.1 |
| Mmp3 | NM_133523.1 |
| Mmp8 | NM_022221.1 |
| Mmp9 | NM_031055.1 |
| Mstn | NM_019151.1 |
| Myd88 | NM_198130.1 |
| Myef2 | NM_001013205.2 |
| Myh6 | NM_017239.2 |
| Myh7 | NM_017240.1 |
| Myh8 | NM_001100485.1 |
| Myl2 | NM_001035252.2 |
| Myl3 | NM_012606.2 |
| Myl4 | NM_001109495.1 |
| Myocd | NM_182667.2 |
| Nf1 | NM_012609.1 |
| Nfatc4 | NM_001107264.1 |
| Nfkb1 | XM_342346.3 |
| Nkg7 | NM_133540.1 |
| Nkx2-5 | NM_053651.1 |
| Nog | NM_012990.1 |
| Nol3 | NM_053516.2 |
| Nos2 | NM_012611.2 |
| Nos3 | NM_021838.2 |
| Notch2 | NM_024358.1 |
| Nr4a1 | NM_024388.2 |
| Nrg1 | NM_001271120.1 |
| Nrp1 | NM_145098.2 |
| Nrp2 | NM_030869.3 |
| Nupr1 | NM_053611.1 |
| Pax2 | NM_001106361.1 |
| Pax3 | NM_053710.1 |
| Pc | NM_012744.2 |
| Pdgfa | NM_012801.1 |
| Pdgfb | NM_031524.1 |
| Pdgfc | NM_031317.1 |
| Pdha1 | NM_001004072.2 |
| Pdk1 | NM_053826.2 |
| Pdk2 | NM_030872.1 |
| Pdk3 | NM_001106581.1 |
| Pdk4 | NM_053551.1 |
| Pecam1 | NM_031591.1 |
| Pecr | NM_133299.1 |
| Pf4 | NM_001007729.1 |
| Pgf | NM_053595.2 |
| Pik3ca | XM_001059296.1 |
| Pik3r1 | NM_013005.1 |
| Pkm | NM_053297.2 |
| Pla2g12a | NM_001108565.1 |
| Pla2g5 | NM_017174.1 |
| POLR1B | NM_031773.1 |
| Ppargc1a | NM_031347.1 |
| Ppargc1b | NM_176075.2 |
| Ppbp | NM_153721.1 |
| Ppp3ca | NM_017041.1 |
| Prkaa1 | NM_019142.1 |
| Prkaa2 | NM_023991.1 |
| Prkca | XM_343975.3 |
| Prkcb | NM_001172305.1 |
| Prkce | NM_017171.1 |
| Prkcg | NM_012628.1 |
| Prl7d1 | NM_053364.1 |
| Prx | NM_023976.2 |
| Ptgs1 | NM_017043.3 |
| Ptk2 | NM_013081.2 |
| Ptn | NM_017066.2 |
| Pxn | NM_001012147.1 |
| Rac1 | NM_134366.1 |
| Rac2 | NM_001008384.1 |
| Raf1 | NM_012639.2 |
| Rheb | NM_013216.1 |
| Rhoa | NM_057132.3 |
| Rnase1 | NM_001029904.1 |
| RPLP0 | NM_022402.2 |
| Runx1 | NM_017325.1 |
| Ryr2 | NM_032078.2 |
| S100b | NM_013191.1 |
| S1pr1 | XM_008761459.1 |
| Sds | NM_053962.3 |
| Serpine1 | NM_012620.1 |
| Shh | NM_017221.1 |
| Ski | XM_017593893.1 |
| Slc2a1 | NM_138827.1 |
| Slc2a4 | NM_012751.1 |
| Slc8a1 | NM_001270772.1 |
| Slc8a2 | NM_078619.1 |
| Slc8a3 | NM_078620.1 |
| Slc9a1 | NM_012652.1 |
| Slit2 | NM_022632.2 |
| Sln | NM_001013247.1 |
| Smad2 | NM_001277450.1 |
| Smad3 | NM_013095.2 |
| Smad4 | NM_019275.2 |
| Smad6 | NM_001109002.2 |
| Smad7 | NM_030858.1 |
| Smo | NM_012807.1 |
| Sod1 | NM_017050.1 |
| Sphk1 | NM_001270811.1 |
| Stat3 | NM_012747.2 |
| Stk11 | NM_001108069.1 |
| Tbx20 | NM_001108132.1 |
| Tbx21 | NM_001107043.1 |
| Tbx5 | NM_001009964.1 |
| Tcap | NM_001271277.1 |
| Tek | NM_001105737.1 |
| Tgfa | NM_012671.2 |
| Tgfb1 | NM_021578.2 |
| Tgfb2 | NM_031131.1 |
| Tgfb3 | NM_013174.2 |
| Tgfbr1 | NM_012775.2 |
| Tgfbr2 | NM_031132.3 |
| Thbs1 | NM_001013062.1 |
| Tie1 | NM_053545.1 |
| Timp1 | NM_053819.1 |
| Timp2 | NM_021989.2 |
| Timp3 | NM_012886.2 |
| Timp4 | NM_001109393.1 |
| Tlr2 | NM_198769.2 |
| Tlr4 | NM_019178.1 |
| Tnfrsf1a | NM_013091.1 |
| Tnnc1 | NM_001034105.1 |
| Tnni3 | NM_017144.1 |
| Tnnt2 | NM_012676.1 |
| Tp53 | NM_030989.3 |
| Tpm1 | NM_001034068.1 |
| Tpm2 | NM_001024345.1 |
| Tpm3 | NM_173111.1 |
| Tpm4 | NM_012678.2 |
| Tpt1 | NM_053867.1 |
| Traf2 | NM_001107815.2 |
| Traf3 | NM_001108724.1 |
| Trx1 | NM_053800.3 |
| Txnrd1 | NM_031614.2 |
| Tymp | NM_001012122.1 |
| Ucn3 | NM_001080208.1 |
| Uqcr10 | NM_001170465.1 |
| Uqcrb | NM_001127553.2 |
| Uqcrc1 | NM_001004250.2 |
| Uqcrc2 | NM_001006970.1 |
| Uqcrfs1 | NM_001008888.1 |
| Uqcrh | NM_001009480.1 |
| Uqcrq | NM_001025134.1 |
| Vcam1 | NM_012889.1 |
| Vegfa | NM_031836.2 |
| Vegfb | NM_053549.1 |
| Vegfc | NM_053653.1 |
| Wars2 | NM_001168641.1 |
| Wt1 | NM_031534.2 |
| Xiap | NM_022231.2 |
| Yap1 | NM_001034002.2 |
